# ViVo: A temporal modeling framework that boosts statistical power and minimizes animal usage

**DOI:** 10.1101/2025.10.14.682266

**Authors:** Guillermo Canudo-Barreras, Eduardo Romanos, Raquel P. Herrera, M. Concepción Gimeno

**Affiliations:** Instituto de Síntesis Química y Catálisis Homogénea (ISQCH) CSIC-Universidad de Zaragoza, C/ Pedro Cerbuna 12, 50009 Zaragoza, Spain; Servicio de Imagen Médica y Fenotipado, Instituto Aragonés de Ciencias de la Salud, Centro de Investigación Biomédica de Aragón (CIBA), Avda. San Juan Bosco, 13, Planta D, E-50009 Zaragoza, Spain

**Keywords:** ViVo platform, Preclinical oncology, Tumor modeling, Tumor Growth Rate (TGR) matrices, Small sample size, 3Rs principles

## Abstract

Preclinical tumor studies are often limited by high variability and small sample sizes, reducing statistical power and masking treatment effects. We present an exponential framework that estimates tumor growth rates (*r*) independently of initial burden and introduces Tumor Growth Rate (TGR) matrices to map treatment effects across time windows. Applied to a public xenograft dataset and four additional mouse models, exponential fits achieved high agreement with raw data (median R^2^ typically > 0.8-0.9). When resampling matched group sizes (n=3-7), our approach consistently outperformed conventional endpoint and daily non-parametric analyses, revealing treatment effects that standard comparisons missed. The framework also predicts tumor weights for animals euthanized early, enabling their inclusion in final analyses and enhancing statistical consistency while advancing the 3Rs principles. To ensure broad adoption, we developed **ViVo**, and open-source web platform (gcanudo-barreras.github.io/ViVo-Platform/) that provides accessible, standardized analysis of *in vivo* tumor kinetics and therapeutic efficacy.

## Introduction

Preclinical oncology faces a critical challenge: while computational advances have revolutionized drug discovery through machine learning^1,2^ and sophisticated algorithms,^3^ practical implementation remains limited by analytical complexity and the need for extensive animal cohorts. Most preclinical studies rely on oversimplified tumor comparisons using classical statistics with insufficient sample sizes, a limitation that contributes to the low success rate, with only 12–36% of compounds showing efficacy in animal models ultimately advancing to Phase III clinical trials.^4^

This analytical gap is evident across recent studies. Meng, Zhang, and coworkers reported critical inconsistencies in which tumor weights revealed significant treatment differences that were not detected by volume measurements (Fig. 1A).^5^ Similar discrepancies were noted by Rong, He, and coworkers, further undermined by statistical flaws such as applying one-way ANOVA to longitudinal tumor growth data (Fig. 1B).^6^ Kolesar and Awuah conducted an efficacy study with only four mice per group, analyzing the data using a Student’s *t*-test, a method with limited robustness for such small sample sizes.^7^ Even bioluminescence imaging faces interpretation difficulties, with different visualization techniques yielding distinct results, thereby compromising data reliability (Fig. 1C).^8^

**Fig. 1:**
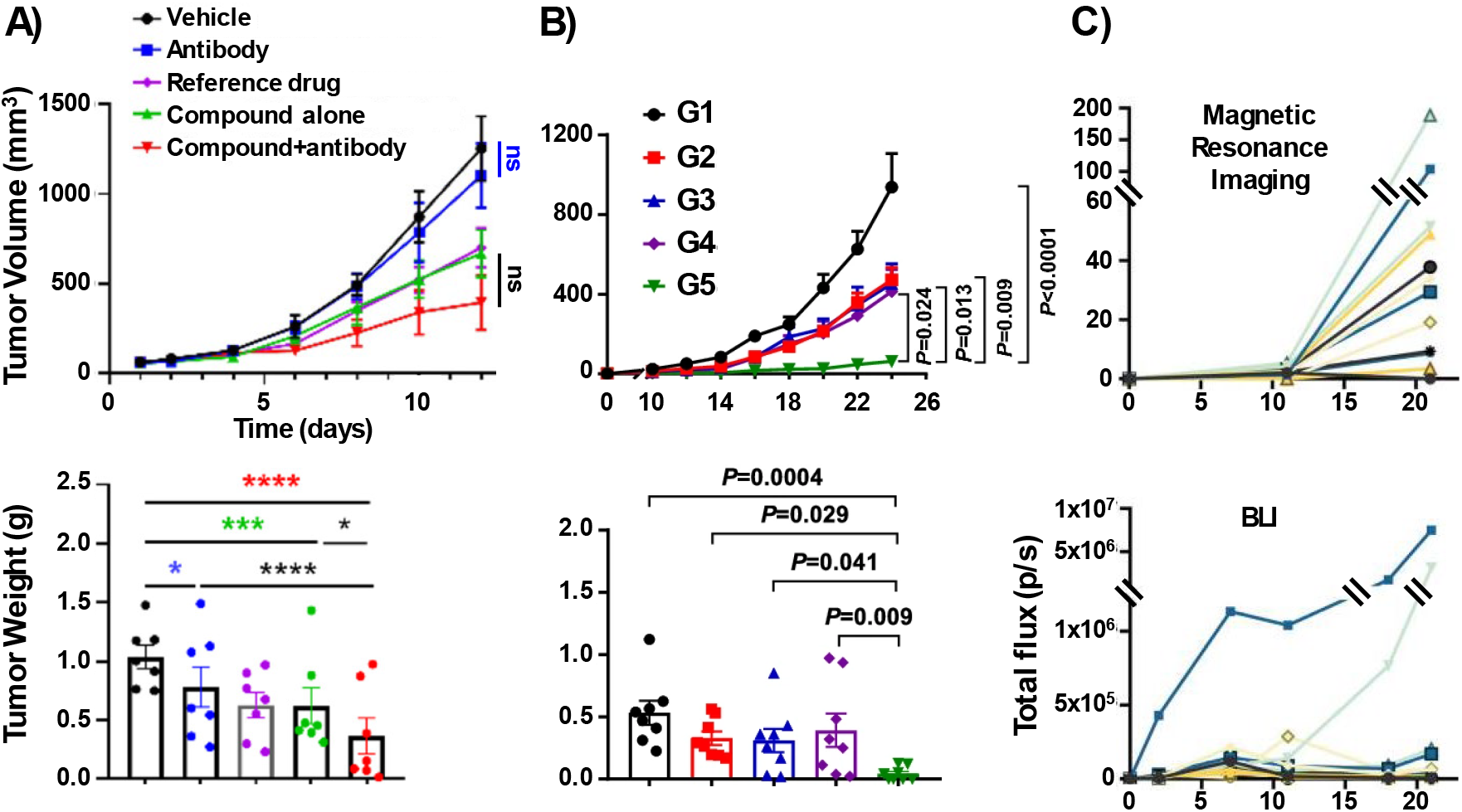
Current methodological limitations in preclinical oncology demonstrate systematic inconsistencies between measurement approaches and inadequate statistical power with small sample sizes. **(A)** ENPP1 inhibitor study (Meng, Zhang) - weight vs volume discrepancies.^5^ **(B)** Polypeptide hydrogel study (Rong, He) - statistical inconsistencies.^6^ **(C)** BLI instability in glioblastoma (Préat, Gallez).^8^

These methodological inconsistencies could theoretically be mitigated by increasing sample sizes, but current research practice is constrained by ethical imperatives that explicitly discourage such approaches.^9^

The internationally recognized 3Rs framework, Replacement, Reduction and Refinement, promotes minimizing animal usage while maximizing data quality.^10^ In parallel, compliance with ARRIVE 2.0 guidelines mandates rigorous study design, transparent reporting, and statistically justified animal numbers.^11^ These principles present a practical and ethical responsibility: to extract richer, more informative insights from smaller cohorts rather than relying on numerical expansion to compensate for analytical limitations. The field requires methodological strategies that enhance interpretability and statistical power without increasing animal burden.

To address this ethical-methodological gap, we propose an exponential modeling solution (Eq. 1):

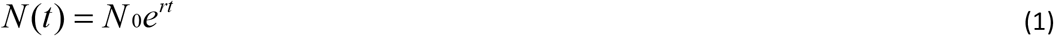

Where *N(t)* represents the number of tumor cells at time *t, N*_*0*_ is the initial tumor burden, and *r* defines the growth rate (time^-1^). This approach enables a fundamental shift from snapshot-based measurements (*N* comparison) to pattern-based analysis (*r* comparison).

The exponential model has proven effective across multiple studies.^12,13^ Both tumor volume and bioluminescence exhibit an approximately linear dependence on cell number, making them suitable parameters for this type of modeling despite their apparent complexity. Two key assumptions underlie the model: cells divide continuously and without constraint. Its application is therefore most appropriate during early-stage tumor growth,^14^ before angiogenesis and nutrient depletion reshape proliferation dynamics.^15^

At the core of our approach are TGR matrices, analytical tools that capture temporal treatment dynamics, exposing growth patterns and trends that remain hidden in endpoint analyses. Beyond descriptive value, time-resolved modeling of tumor size provides predictive capabilities, such as estimating tumor weight on defined days relative to sacrifice. This enables standardized tumor weight comparisons across animals within a study, even when euthanasia occurs at different times due to humane endpoint criteria.

These methodologies are implemented in the ViVo platform, an accessible, web-based tool for systematic tumor growth analysis (Fig. 2). Beyond enabling exponential modeling, TGR matrices, and time-resolved tumor weight standardization, ViVo also facilitates the systematic identification and removal of outliers and includes an advisory tool to assess animal model homogeneity, providing researchers with a standardized approach to improve data consistency and interpretability without increasing cohort sizes. Together, these features allow more reliable, ethically responsible analysis of preclinical tumor studies, extracting richer insights from limited animal datasets.

**Fig. 2:**
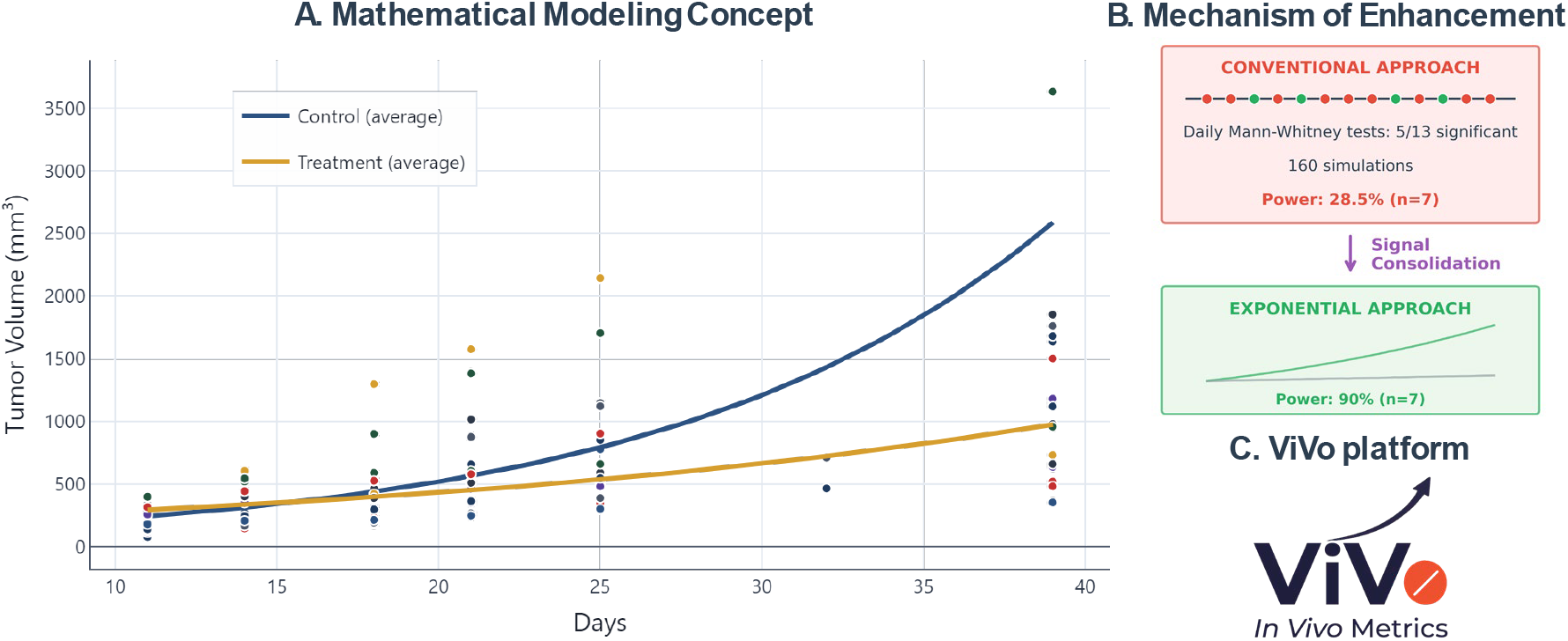
Schematic representation of the ViVo platform, which applies exponential modeling and temporal analysis to convert conventional tumor measurements into robust growth parameters.

## Results

### Enhanced Statistical Power with Reduced Sample Size Requirements

Exponential modeling demonstrated superior statistical power across all tested sample sizes, maintaining performance levels unattainable through conventional analysis. To confirm model applicability and assess its statistical power, we used the publicly available dataset of Dr. Constantine Daskalakis (https://www.causeweb.org/tshs/tumor-growth/), derived from a subcutaneous xenograft mouse model with human glioma cells consisting of 18 animals (8 control and 10 treated mice, n=8,10) with 13 timepoints (days 0, 3, 4, 5, 6, 7,10, 11, 12, 13, 14, 17, and 18).^16^

Mathematical modeling achieved exceptional goodness-of-fit across the dataset. 17/18 animals achieved coefficient of determination R^2^ > 0.8 (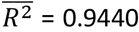 for control group, 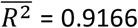 for drug group), while 13/18 reached R^2^ > 0.9 (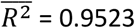 for control group, 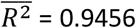 for drug group). With R^2^ > 0.8 filtering, exponential modeling revealed a 31.9% reduction in growth rate between treated and control groups (*r* = 0.175 ± 6.9% vs 0.257 ± 11.3% day^−1^, respectively). Analogously, R^2^ > 0.9 filtering achieved a 29.8% reduction in growth (*r* = 0.193 ± 5.8% vs 0.274 ± 10.8% day^−1^; Supplementary Table S1).

Systematic evaluation across eight sample size scenarios (from n=10,8 down to n=3,3 *per* group) revealed the exceptional performance of the framework in limited-sample contexts (Table 1).

**Table 1.** Statistical Power Comparison Using Daskalakis Public Dataset.^16^.

| Entry | Sample Size<br>*n* per group | Conventional Analysis<br>Statistical Power (%) | Exponential Model |  | Improvement<br>vs Conventional |
| --- | --- | --- | --- | --- | --- |
| | | | $R^2 > 0.8$ | $R^2 > 0.9$ | |
| 1 | $n=10, n=8$ | 61.5 | 100 | 100 | +38.5% |
| 2 | $n=9, n=8$ | 53.1 | 100 | 100 | +46.9% |
| 3 | $n=8, n=8$ | 41.9 | 100 | 100 | +58.1% |
| 4 | $n=7, n=7$ | 28.5 | 40 | 90 | +61.5% |
| 5 | $n=6, n=6$ | 22.3 | 30 | 70 | +47.7% |
| 6 | $n=5, n=5$ | 9.7 | 40 | 40 | +30.3% |
| 7 | $n=4, n=4$ | 7.0 | 20 | 35* | +28% |
| 8 | $n=3, n=3$ | 0.0 | 35 | 40* | +40% |

\*For extremely small samples ( $n \leq 4$ ), subsets with insufficient animals after quality filtering were excluded from power calculations.

Mathematical modeling demonstrated superior statistical power across all configurations, maintaining 100% performance down to n=8,8 groups (Table 1, entries 1-3) and 90% power with n=7,7 samples (Table 1, entry 4). Even at extreme sample sizes (n=3,3), exponential modeling achieved 35-40% power, compared to 0% for conventional analysis (Table 1, entry 8). Conventional analysis employed daily Mann-Whitney U tests with statistical power calculated as the percentage of days achieving significance, reaching a maximum of 61.5% even with the largest cohort tested (Table 1, entry 1).

This enhancement is visualized across the complete sample size range (Fig. 3), where exponential modeling maintains robust performance even with extremely limited cohorts.

**Fig. 3:**
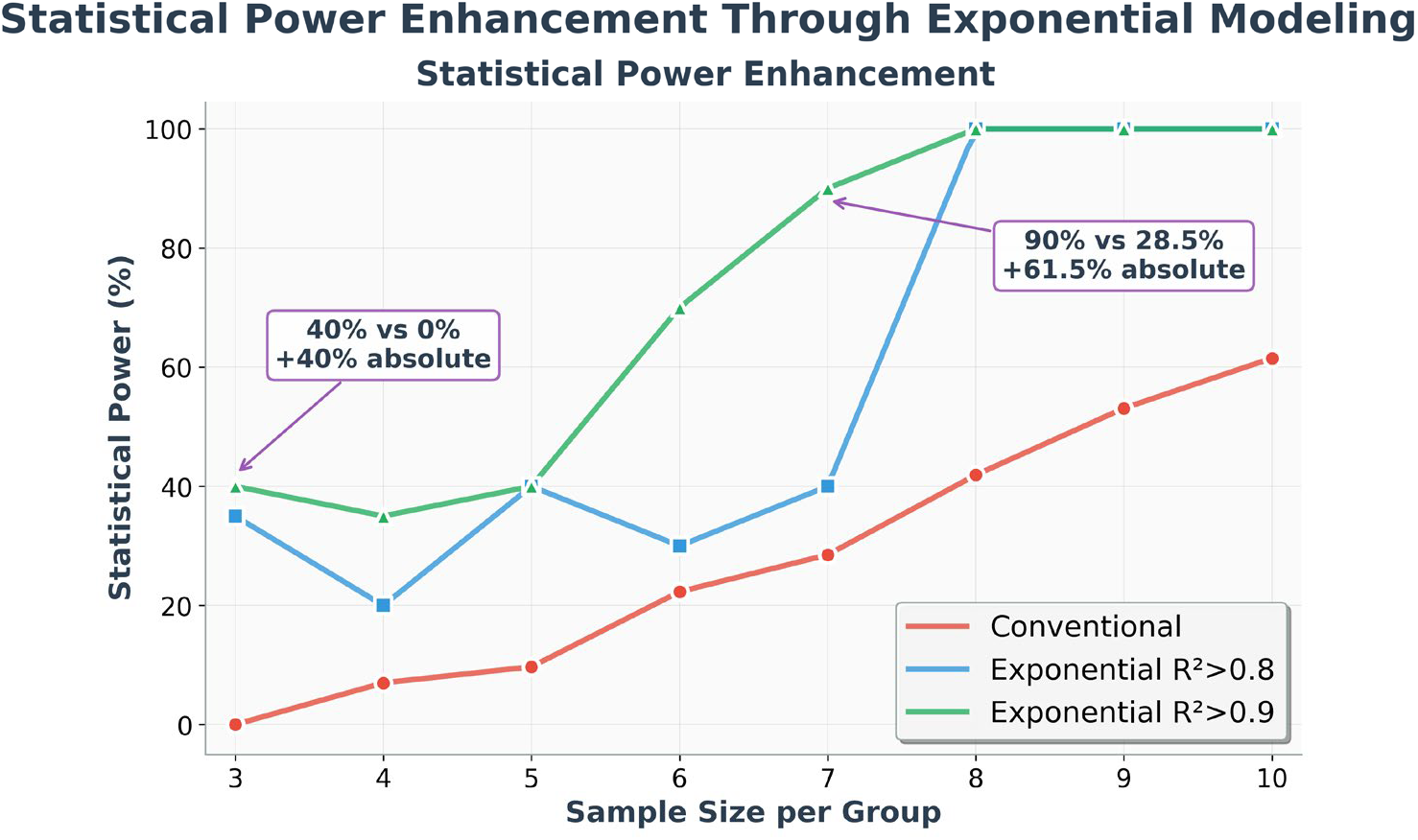
Exponential modeling enhances statistical power across all sample sizes. Statistical power comparison using Daskalakis dataset: conventional analysis (red circles), exponential modeling with quality filters R^2^ > 0.8 (blue squares) and R^2^ > 0.9 (green triangles).

The mechanistic basis for this improvement arises from fundamental principles of error propagation (Supplementary Note 1). Analysis of parameter variability demonstrates a systematic advantage of evaluating growth rates over tumor volume measurements. Specifically, the relative standard error (RSE) for growth rates is consistently lower: RSE(*r*) = 9.1% versus RSE(*V*_*0*_) = 17.6%, corresponding to a 48% reduction in parameter uncertainty. In exponential growth models, uncertainty in the growth rate scales quadratically with time, whereas uncertainty in the initial tumor burden contributes linearly to measurement variability (Eq. 2).

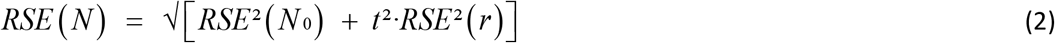

Consequently, using growth rates instead of raw biomarker data for comparison reduces variability while enabling robust long-term analysis.

### Temporal Pattern Discovery through TGR Matrix Analysis

While exponential modeling demonstrated superior statistical power by extracting single growth parameters from entire longitudinal studies, systematic application of Tumor Growth Rate matrices revealed that treatment efficacy exhibits distinct temporal heterogeneity invisible to conventional endpoint analysis. Real therapeutic interventions often exhibit temporal dynamics: drug bioavailability fluctuates, resistance mechanisms emerge, and cellular vulnerabilities shift over time; creating treatment dynamics that single-parameter models cannot detect.

To capture this temporal complexity while keeping mathematical rigor, we extended the exponential modeling principle to localized temporal intervals. Rather than calculating one growth rate for the entire study period, we systematically calculated growth rates between two timepoints x and y (Eq. 3):

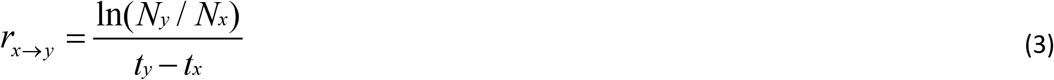

This localized growth rate (*r*_*x*→*y*_) represents the instantaneous growth dynamics between specific timepoints.

When applied to the Daskalakis dataset, TGR analysis generated 78 distinct temporal intervals (all possible day-to-day combinations; Fig. 4A-B), compared to 13 single timepoints in conventional analysis, providing 6-fold higher resolution. This enhanced resolution revealed three main tumor growth phases with distinct therapeutic characteristics.

**Fig. 4:**
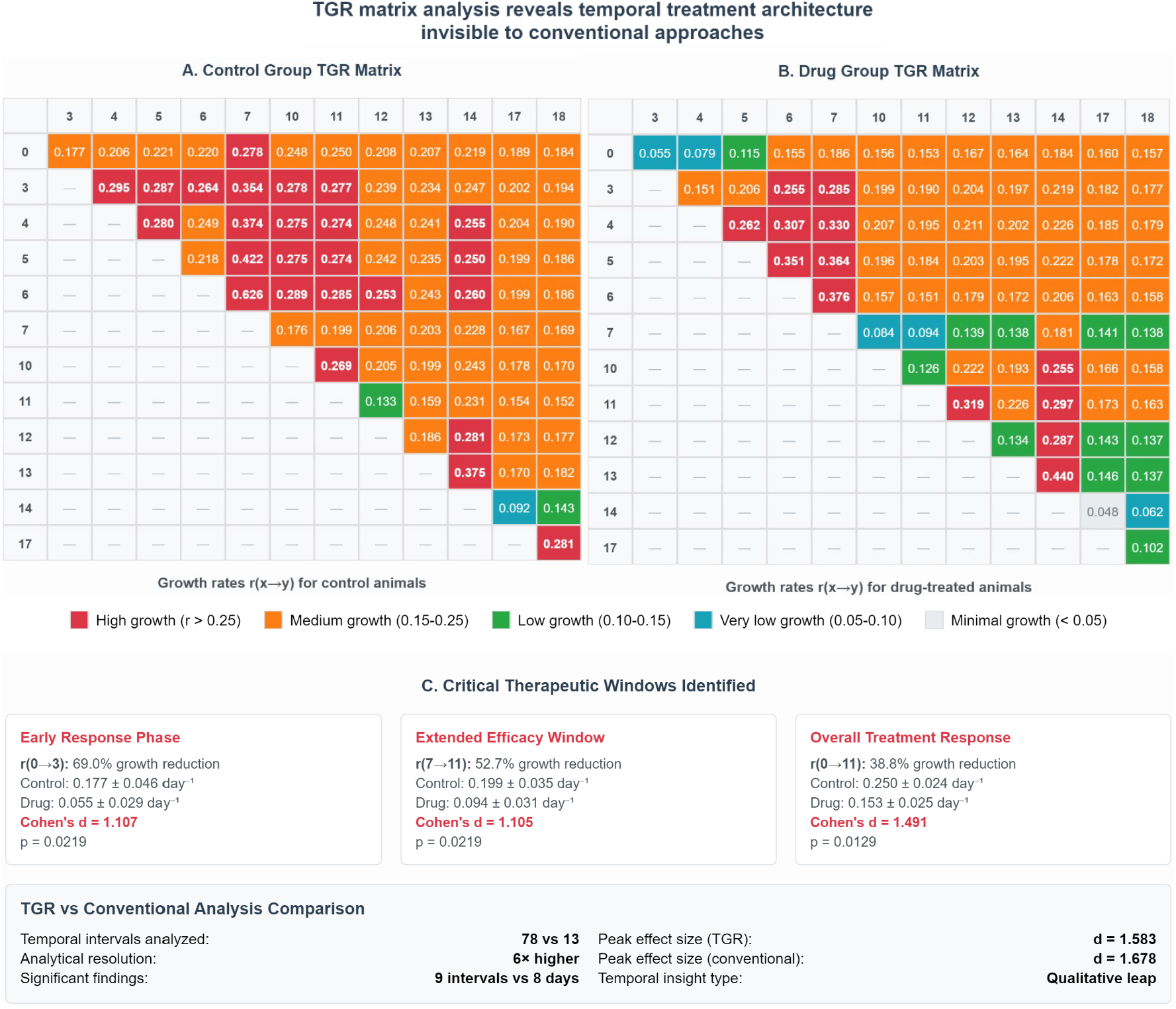
Temporal Tumor Growth Rate (TGR) Matrix Analysis. **(A-B)** TGR matrices for Control **(A)** and Drug-treated **(B)** groups showing growth rates (day^−1^) between all timepoint pairs from day 0 to 18. Color intensity reflects growth rate magnitude: red indicates high growth rates, orange indicates medium rates, green indicates low rates, blue indicates very low rates, and pale gray indicates minimal growth, zero or negative rates. **(C)** Statistical summary of the three tumor growth phases identified through systematic TGR matrix analysis.

<u>Early Response</u> (*r*_*0*→*3*_) demonstrated the most pronounced effect, with 69.0% growth rate reduction observed in the first three days (0.055 ± 0.029 vs 0.177 ± 0.046 day^-1^ control, p-value =0.0219, Cohen’s d=1.107).

<u>Extended Efficacy</u> (*r*_*7*→*11*_) showed 52.7% sustained suppression during mid-treatment phase (0.094 ± 0.031 vs 0.199 ± 0.035 day^-1^ control, p-value =0.0219, Cohen’s d=1.105), from day 7 to 11.

<u>Long-Term Response</u> (*r*_*0*→*11*_) maintained 38.8% comprehensive efficacy (0.153 ± 0.025 vs 0.250 ± 0.024 day^-1^ control, p-value =0.0129, Cohen’s d=1.491) after 11 days of treatment.

While traditional statistical comparisons identified differences on 8 of 13 study days (Supplementary Table S2), TGR analysis identified significant treatment effects in 9 of 78 intervals (11.5%) with effect sizes ranging from Cohen’s d = 1.105 to d= 1.583 (Supplementary Note 2). Multi-day intervals consistently achieved large effect sizes (Cohen’s d > 1.1), with peak efficacy during the *r*_*0*→*3*_ window, followed by the *r*_*7*→*11*_ phase. These systematic temporal patterns demonstrate heterogeneity that is invisible to single-day measurements and undetectable by traditional statistical approaches (Fig. 4C), uncovering therapeutic dynamics that could guide dosing schedules and the design of combination therapies.

### Cross-Platform Validation and Predictive Applications

The framework demonstrated robust performance across four additional experimental models: three orthotopic tumor models in athymic nude mice using both human and murine cells lines with different kinetic profiles, plus one subcutaneous syngeneic model in immunocompetent mice. Animal models included: **(1)** syngeneic orthotopic mammary gland adenocarcinoma using 4T-1-Luc2 cells in 16 athymic mice (n=8,8); **(2)** xenogeneic orthotopic triple-negative breast cancer using MDA-MB-231-luc2-GFP human cells in 12 athymic nude mice (n=6,6); **(3)** syngeneic orthotopic pancreatic ductal adenocarcinoma (PDAC) using MLK 3287-Luc cells in 16 athymic nude mice (n=8,8); and **(4)** subcutaneous syngeneic breast model using 4T-1-Luc2 cells in 21 immunocompetent BALB/cByJRjmice (n=10,11), with data from our previous study “Synthetic ease and exceptional *in vivo* performance of pyrazole-based cyclometallated iridium complexes”.^17^

After applying R^2^ > 0.8 filtering, mathematical modeling proved successful across all experimental groups, with R^2^ values between 0.8683 and 0.9586, and valid animal percentages above 67% in all cases (Table 2).

**Table 2.** Exponential fitting metrics to the used mouse models data.

| Entry | Mouse Model | Group | $N_0$ | $r$ (day<sup>-1</sup>) | $R^2$ | Valid Animals |
| --- | --- | --- | --- | --- | --- | --- |
| 1 | Orthotopic 4T-1-Luc2<sup>*1</sup> | Control | 2.99E+8±**10.6%** | 0.670±**4.7%** | **0.9357** | 7/8 |
| 2 |  | Treatment | 2.94E+8±**24.9%** | 0.534±**9.1%** | **0.9192** | 6/8 |
| 3 | Xenogeneic MDA-MB-231-luc2-GFP<sup>*2</sup> | Control<sup>*3</sup> | 5.4±**60.1%** | 0.135±**19.7%** | **0.8683** | 4/6 |
| 4 |  | Treatment | 4.9±**43.4%** | 0.146±**13.0%** | **0.9500** | 6/6 |
| 5 | Orthotopic MLK 3287-Luc<sup>*1</sup> | Control | 7.69E+7±**37.6%** | 0.232±**8.4%** | **0.9092** | 7/10 |
| 6 |  | Treatment | 1.05E+8±**24.4%** | 0.177±**8.9%** | **0.8928** | 8/10 |
| 7 | Subcutaneous 4T-1-Luc2<sup>*2</sup> | Control | 74.2±**17.9%** | 0.107±**6.1%** | **0.9586** | 10/10 |
| 8 |  | Treatment | 152.0±**16.6%** | 0.054±**11.9%** | **0.9475** | 8/11 |

<sup>\*1</sup> Tumor size was estimated by BLI, then $N_0 = BLI_0$ <sup>\*2</sup> Tumor size was estimated by caliper, then $N_0 = V_0$ (mm<sup>3</sup>) <sup>\*3</sup> $R^2$ threshold lowered to > 0.7 due to extremely small sample size (n=3 valid animals with $R^2 > 0.8$ ).

Relative standard errors for initial tumor burden were consistently higher than those for growth rates (RSE(*N*_*0*_) > RSE(*r*)). Growth rates analysis revealed substantial kinetic differences between cell lines: 4T-1-Luc2 cells exhibited *r* = 0.670 day^-1^ compared to *r* = 0.135 day^-1^ for MDA-MB-231-luc2-GFP cells.

Once mathematical modeling proved successful in describing all models, growth rates were compared between experimental groups. The results demonstrated both concordance with conventional analysis and enhanced detection capabilities:

#### Positive Control Validation

Traditional statistical comparison of orthotopic 4T-1-Luc2 model data (Supplementary Table S3) revealed significant differences on days 4 (p-value = 0.015) and 7 (p-value < 0.001), with corresponding growth rate differences (p-value = 0.012, Cohen’s d = 1.341). Conventional statistical analysis of subcutaneous 4T-1-Luc2 model data (Supplementary Table S4) also showed significant differences on days 21 (p-value = 0.045, Cohen’s d = 1.194) and 32 (p-value = 0.005, Cohen’s d = 1.381), with tumor growth differences (p-value < 0.001, Cohen’s d = 2.736).

#### Negative Control Validation

No significant treatment effects were observed for the xenogeneic orthotopic MDA-MB-231-luc2-GFP mouse model in either conventional analysis (Supplementary Table S5) or growth rate comparisons (p-value = 0.366, Cohen’s d = 0.348).

#### Enhanced Detection Capability

While conventional analysis detected no significant differences across study duration for the orthotopic MLK 3287-Luc animal model (Supplementary Table S6), mathematical modeling and tumor growth comparison revealed significant rate differences (p-value = 0.043, Cohen’s d = 1.146).

TGR analysis of each model revealed distinct efficacy phases across the different studies:

#### Syngeneic Orthotopic 4T-1-Luc2

The control group maintained sustained high growth rates throughout the study, while the treatment group showed significantly lower growth rates from day 4 to 7 (*r*_*4*→*7*_: 0.502 vs 0.756 day^-1^ control; Fig. 5A), representing a 33.6% growth reduction (p-value = 0.007, Cohen’s d = 1.471).

**Fig. 5:**
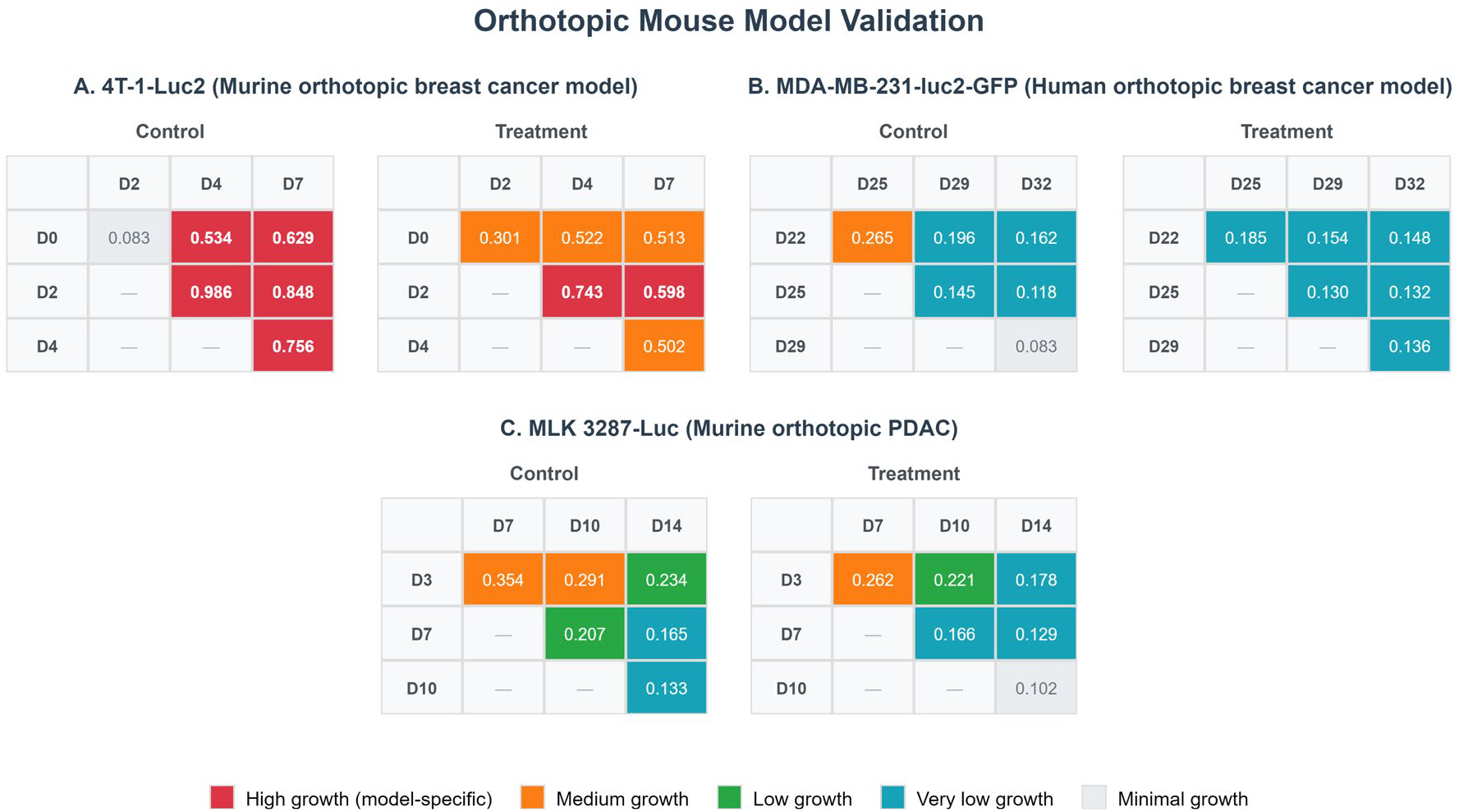
Validation of orthotopic mouse models through TGR analysis. **(A)** TGR matrices for the murine breast 4T-1-Luc2 model (days 0-7). **(B)** TGR matrices for the human triple-negative breast MDA-MB-231-luc2-GFP model (days 22-25). **(C)** TGR matrices for the murine PDAC MLK 3287-Luc model (days 3-14).

#### Xenogeneic Orthotopic MDA-MB-231-luc2-GFP

Both groups exhibited similar growth rates across all TGR phases, with no significant differences between control and treatment groups in any interval (Fig. 5B).

#### Syngeneic Orthotopic MLK 3287-Luc

Control and treatment groups showed longitudinal deceleration over time (control: *r*_*3*→*7*_ = 0.354 day^-1^ → *r*_*10*→*14*_ = 0.133 day^-1^; treatment: *r*_*3*→*7*_ = 0.282 day^-1^ → *r*_*10*→*14*_ = 0.102 day^-1^), evidencing a distinct kinetic profile compared to previous models. Nonetheless, the treatment group experienced greater decline with significant differences emerging in the final study interval (*r*_*3*→*14*_: 0.178 vs 0.234 day^-1^ control; p-value = 0.049; Cohen’s d = 1.245; Fig. 5C).

#### Subcutaneous Syngeneic 4T-1-Luc2

From day 11, the treated group maintained consistently lower growth rates than control, with peak 67.8% reduction from day 14 to 21 (*r*_*14*→*21*_: 0.039 vs 0.122 day^-1^ control). Only one animal in the control group reached day 39. Hence, statistical comparisons between TGR intervals that involved day 39 were not possible. For the 15 valid time intervals, significant differences were observed for 12 of them (Supplementary Table S7).

Mathematical modeling enabled accurate prediction of tumor weights across different euthanasia dates, addressing the practical challenge of humane endpoint criteria. Exponential fitting for the subcutaneous 4T-1-Luc2 model yielded 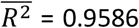 for control group and 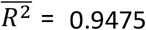 for treated group (Table 2, entries 7 and 8). Direct validation on day 39 was possible for five animals where both predicted and experimental tumor weights were available. The comparison between experimental and predicted values demonstrated prediction errors ranging from 1.2% to 40.6%, with most estimates within 11% accuracy (Table 3).

**Table 3.** Tumor Weight Estimations on day 39 with caliper-based measurement-prediction deviation.

| Entry | Group | Euthanasia date | Experimental Weight (g) | Predicted Weight (g) | Error (%)* |
| --- | --- | --- | --- | --- | --- |
| 1 | Control | 32 | 0.993 | 0.400 | - |
| 2 | Control | 32 | 1.167 | 0.500 | - |
| 3 | Control | 32 | 1.742 | 0.700 | - |
| 4 | Control | 32 | 2.14 | 0.900 | - |
| 5 | Control | 39 | 0.634 | 0.600 | 5.6 |
| 6 | Control | 25 | 6.004 | 0.900 | - |
| 7 | Control | 32 | 0.605 | 0.300 | - |
| 8 | Control | 32 | 1.133 | 0.400 | - |
| 9 | Control | 32 | 1.375 | 0.600 | - |
| 10 | Control | 32 | 1.554 | 0.600 | - |
| 11 | Treatment | 32 | 0.798 | 0.500 | - |
| 12 | Treatment | 39 | 0.311 | 0.350 | 11.1 |
| 13 | Treatment | 32 | 0.631 | 0.400 | - |
| 14 | Treatment | 39 | 0.395 | 0.400 | 1.2 |
| 15 | Treatment | 39 | 0.703 | 0.500 | 40.6 |
| 16 | Treatment | 39 | 0.328 | 0.300 | 9.4 |
| 17 | Treatment | 32 | 0.495 | 0.400 | - |
| 18 | Treatment | 32 | 0.638 | 0.400 | - |

\* Prediction error was calculated as the difference between experimental and predicted caliper measurements related to the first.

Predicted tumor weights on day 39 were 1.40 ± 0.59 g (mean ± SD) for controls and 0.52 ± 0.19 g for the treatment group, with hybrid experimental-predicted groups showing significant differences (p-value = 0.003, Cohen’s d = 1.069). This predictive capability facilitates standardization of tumor weights across all animals within the same study, even when euthanasia occurs on different dates due to humane endpoint criteria.

These analytical approaches are implemented in the freely available ViVo web platform (<u>gcanudo-barreras.github.io/ViVo-Platform/</u>), providing accessible computational workflows that enable direct application of exponential modeling, TGR matrix analysis and tumor weight predictive and standardization capabilities without specialized expertise requirements.

## Discussion

The ViVo framework demonstrates that mathematical modeling can substantially enhance statistical power while reducing animal requirements in preclinical oncology. The mechanistic advantage becomes particularly pronounced in small sample scenarios, where conventional analyses often fail to detect genuine treatment effects. Our systematic evaluation reveals that exponential modeling maintains superior statistical performance across all tested sample sizes, achieving 90% power with n=7,7 groups, and 100% power above, compared to 61.5% maximum for conventional analysis with larger cohorts. This enhancement enables approximately 40% reduction in animal usage while maintaining robust detection capabilities, as demonstrated by the Daskalakis dataset analysis where comparable statistical outcomes required substantially fewer animals.

The introduction of Tumor Growth Rate (TGR) matrices represents a methodological advance, transforming discrete measurements into comprehensive temporal maps of treatment dynamics. Our analysis revealed three distinct therapeutic phases: Early Response, Extended Efficacy, and Long-Term Response, that remained invisible to conventional endpoint analysis. Notably, the observed kinetic differences between cell lines align with established literature, with 4T1 cells characterized as rapidly proliferating (https://www.atcc.org/products/crl-2539-luc2)^18^ and MDA-MB-231 cells as comparatively indolent (https://www.atcc.org/products/htb-26).^19^ The resulting 5-fold quantitative differentiation validates that the framework captures genuine biological kinetics rather than mathematical artifacts, providing confidence in the ability of the method to discriminate meaningful therapeutic effects from experimental noise.

This temporal granularity provides quantitative insights into treatment kinetics, revealing that therapies with similar overall efficacy may exhibit markedly different response patterns. The systematic identification of peak efficacy windows has direct implications for optimizing dosing schedules and designing rational combination therapies based on drug action kinetics.

### Methodological Positioning and Applicability

Compared to alternative mathematical approaches, exponential modeling offers an optimal balance between biological relevance and practical applicability for early-stage tumor dynamics. While logistic and Gompertz models provide theoretical advantages for late-stage progression, they require extensive datasets and prolonged observation periods to capture critical parameters such as carrying capacity and inflection points.^1,20^ These requirements often conflict with standard preclinical protocols,^21,22^ which typically span 2-4 weeks and focus on proliferation-dominated phases where exponential assumptions remain valid.^2,11^

The exponential framework becomes particularly advantageous compared with modern AI-based approaches, which rely heavily on large, high-quality datasets for pattern generalization.^23^ In sample-limited settings, parameter-rich models suffer from reduced identifiability and compromised robustness.^24^ Our approach addresses this challenge by extracting biologically meaningful parameters from limited data while maintaining mathematical rigor through goodness-of-fit thresholds and systematic quality control.

A critical aspect of successful implementation is recognizing appropriate application boundaries. The exponential model accurately describes tumor dynamics during proliferation-dominated phases but becomes less reliable when angiogenic constraints and nutrient depletion alter growth kinetics. Objective indicators of model breakdown, including systematic residual patterns and R^2^ values falling below 0.8, serve as quality control thresholds, ensuring analytical reliability within well-defined operating ranges.

### Limitations and Critical Assessment

Several important limitations should be acknowledged when implementing this framework. The exponential growth assumption fundamentally restricts applicability to early-stage tumor development, typically within the first 2-4 weeks of preclinical studies. Beyond this window, more complex growth dynamics emerge that require alternative modeling approaches. Additionally, the quality filtering process, while necessary for parameter reliability, can exclude animals with poor model fits, potentially introducing selection bias that may not reflect true treatment responses.

Mathematical modeling faces intrinsic statistical limitations when applied to extremely small groups (n ≤ 4). Even with optimal model performance, such samples sizes constrain the reliability of parameter estimates due to fundamental statistical principles rather than methodological limitations. While our approach improves robustness in limited-sample contexts, thoughtful experimental design remains essential for ensuring valid and interpretable results.

The temporal analysis provided by TGR matrices, while offering enhanced resolution, requires careful interpretation to avoid false discoveries arising from multiple comparisons. Although we systematically evaluated 78 temporal intervals compared to 13 conventional timepoints, this increased analytical resolution must be balanced against appropriate statistical corrections to maintain Type I error control. Furthermore, the biological interpretation of short-term growth rate fluctuations requires careful consideration of measurement precision and potential artifacts.

### Impact and Practical Applications

The alignment of this methodology with 3Rs principles represents perhaps its most significant contribution to preclinical research. Our demonstration that mathematical modeling can achieve superior statistical performance with smaller sample sizes suggests potential for meaningful animal reduction across preclinical oncology. Analysis of the Daskalakis dataset indicates that exponential modeling achieves 70% statistical power using 12 animals while conventional analysis maintains 61.5% power with 18 animals, representing nearly 40% reduction in animal usage while improving analytical precision.

TGR analysis provides a promising tool for early detection of therapeutic response, determination of optimal treatment windows, and early resistance signals. The systematic identification and removal of outliers, combined with standardized data quality assessment, addresses a critical challenge in preclinical studies where inconsistent data handling can compromise reproducibility. The predictive capabilities demonstrated through tumor weight standardization offer immediate practical benefits for addressing humane endpoint requirements. This application alone may justify implementation in laboratories struggling with data interpretation across heterogenous sacrifice dates. The freely available ViVo platform embodies these principles, providing automated quality control and interactive analysis tools that are accessible to researchers without computational expertise.

## Conclusions

This framework demonstrates that mathematical modeling can substantially enhance statistical power in preclinical tumor studies while reducing animal usage by up to 40%. By converting conventional measurements into growth rate parameters and enabling temporal pattern analysis through TGR matrices, the approach addresses fundamental challenges in small-sample preclinical research where biological variability often masks genuine treatment effects.

The structured temporal data foundation provided by TGR matrices creates natural integration points for machine learning applications in precision oncology, positioning the framework as a potential bridge between conventional preclinical assessment and emerging computational approaches in personalized cancer therapy. As precision medicine strategies gain prominence in clinical oncology, the need for standardized preclinical evaluation methodologies that extract maximum information from limited samples becomes increasingly critical.

While our current validation encompasses multiple cell lines and implantation sites, broader validation across diverse tumor models, particularly patient-derived xenograft (PDX) systems, remains essential for establishing methodological robustness. Several limitations should be acknowledged, including the exponential growth assumption that may not capture all tumor growth dynamics, and the R^2^ thresholds, which should be adjusted judiciously to avoid introducing selection bias. Integration with existing laboratory information management systems and regulatory submission pathways will facilitate adoption across research institutions and pharmaceutical development programs.

The ViVo framework ultimately represents a convergence of mathematical rigor, practical accessibility, and ethical responsibility in preclinical research. By enhancing analytical precision while reducing animal usage, this approach establishes a foundation for more efficient and responsible preclinical oncology research.

## Experimental Section

### Mathematical Framework

#### Exponential Growth Modeling

Growth rate parameters (*r*) were extracted through linear regression on log-transformed data. Quality assessment used coefficient of determination (R^2^) with configurable thresholds. We set R^2^> 0.8 as the default quality threshold, following established practice in tumor growth modeling where exponential models typically achieve R^2^ values exceeding 0.8-0.9 when applied to appropriate early-stage data. This threshold balances model selectivity with practical applicability, ensuring reliable parameter extraction while avoiding overly restrictive filtering that could exclude biologically meaningful data.

Tumor volume was estimated using the standard ellipsoid formula (Eq. 4):

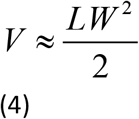

Where L and W represent length and width measurements respectively.

#### Model Assumptions and Validity Criteria

Model validity requires exponential growth conditions typically present during the first 2-4 weeks of preclinical studies. Application boundaries are defined by multiple criteria:

- R^2^ threshold filtering (default >0.8, configurable 0.0-1.0)
- Residual pattern analysis for systematic deviations from exponential behavior
- Temporal applicability boundaries based on study duration and growth phase
- Automatic flagging when growth patterns suggest model breakdown (systematic residual patterns, R^2^ degradation below 0.8)

Indicators of model breakdown serve as objective boundaries for valid application, ensuring analytical reliability within well-defined operating ranges compatible with standard efficacy assessment protocols.

#### Tumor Growth Rate (TGR) Matrix construction

For each temporal analysis, localized growth rates were calculated between all possible timepoint pairs using Eq. 3. This localized growth rate (*r*_*x*→*y*_) represents the instantaneous growth dynamics between specific timepoints x and y, enabling detection of temporal patterns invisible to whole-study analysis. When calculated for all possible time combinations, the localized growth rates form structured matrices where each element *r*_*x*→*y*_ describes a specific temporal interval, with rows corresponding to starting time points and columns to ending time points.

### Automated Quality Control System

Before fitting the data into the exponential model, the Automated Quality Control System evaluates data homogeneity, searches for potential outliers and analyzes them. The system employs interquartile range (IQR) analysis with configurable sensitivity thresholds that can be adjusted both automatically—based on sample size—or manually. The <u>three sensitivity levels</u> are:

- Ultra-Conservative (n <8)
- Conservative (n = 8-12)
- Moderate (n > 12)

#### Homogeneity Assessment

Before analysis, ViVo ensures data homogeneity using coefficient of variation (CV) and a Homogeneity Quality Score that accounts for both CV and sample size. Assessment occurs twice: before and after outlier filtering to ensure transparency in data processing decisions.

For each animal, the baseline value is the first recorded measurement, with non-positive baseline values excluded. The Homogeneity Quality Score combines a piecewise-linear penalty on CV with multiplicative corrections for small sample sizes. The base score uses two thresholds (ε = 15 for excellent, π = 30 for poor), with the formula (Eq. 5):

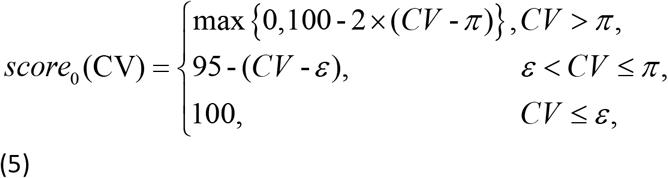

A multiplicative adjustment factor accounts for small sample uncertainty (Eq. 6):

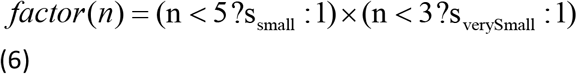

With the final score is computed as (Eq. 7)

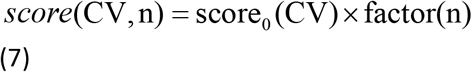

#### Six-Criteria Outlier Detector

The platform implements systematic outlier detection through six criteria with severity-based classification (Supplementary Table S9). The system employs interquartile range (IQR) analysis on log-transformed data with configurable sensitivity that automatically adjusts based on sample size:

##### Critical Severity

- IMPOSSIBLE_VALUE: Values ≤ 0
- EXTREME_GROWTH: Excessive exponential growth (>1000-5000%/day depending on sample size)
- EXTREME_DECLINE: Excessive exponential decline (>67-90%/day depending on sample size)

##### High Severity

- INTRA_OUTLIER: Outliers within individual animals using IQR analysis

##### Medium Severity

- GROUP_OUTLIER: Outliers compared to group median
- LAST_DAY_DROP: Abrupt drops on final measurement day

Sensitivity automatically adjusts with three levels: Ultra-Conservative (n<5, 4.0× IQR multiplier), Conservative (n=5-10, 3.0× multiplier), and Moderate (n>10, 2.0× multiplier). For small samples, multiple indicators are required for outlier classification to prevent excessive data exclusion (Supplementary Table S9).

#### Dual Analysis Framework

Three complementary analysis approaches accommodate different data quality scenarios:

1. <u>Complete Analysis</u>: All animals and measurements included regardless of quality flags
2. <u>Animal-Level Filtering</u>: Entire animals excluded when containing selected anomalies
3. <u>Point-Level Filtering</u>: Individual problematic measurements excluded while retaining animals with ≥3 valid timepoints

This framework enables transparent assessment of outlier impact on study conclusions while keeping analytical rigor, allowing researchers to evaluate how quality control decisions affect their results.

### Statistical Analysis and Interactive Framework

#### Primary Statistical Comparisons

Growth rate comparisons between experimental groups employed Mann-Whitney U tests to handle small sample sized and avoid normality assumptions inherent to limited sample sizes.

#### Power Assessment Methodology

Statistical power evaluation employed systematic random subset selection from the original Daskalakis dataset (n=8,10). For each sample size scenario (n=8,10 down to n=3,3), 20 random subsets were generated to account for selection variability. For conventional analysis, statistical power was calculated as the percentage of study days significance (p-value < 0.05) averaged across all subsets with each scenario. For exponential modeling, power was calculated as the proportion of subsets detecting significant growth rate differences, with results categorized as binary outcomes (significant/non-significant) for each subset.

#### TGR Matrix Statistical Implementation

For TGR analysis, statistical comparisons utilize complete individual animal data while displayed matrices show group-averaged values for visualization. The interactive analysis framework enables real-time statistical testing of specific temporal intervals through point-and-click selection, with automatic calculation of:

- Group medians and interquartile ranges for each temporal interval
- Mann-Whitney U test statistics and p-values for between-group comparisons
- Cohen’s d effect sizes with 95% confidence intervals

Effect sizes follow Cohen’s conventions: small (d=0.2-0.5), medium (d=0.5-0.8), and large (d≥0.8), with comprehensive outputs including test statistics, significance determinations, and confidence intervals formatted for publication.

#### Variance Reduction Quantification

Parameter variability was assessed using relative standard error (RSE). The variance reduction mechanism follows error propagation principles (Supplementary Note 1) where growth rate uncertainty scales quadratically with time (Eq. 2).

### Platform Implementation and Technical Specifications

#### System Requirements and Compatibility

The ViVo platform operates entirely in-browser using JavaScript ES6+, requiring no software installation or server-side processing. Compatible browsers include Chrome 60+, Firefox 55+, Safari 12+, and Edge 79+. Modern browser compatibility ensures broad accessibility across research environments while supporting FileReader API and CSS Grid for optimal functionality.

#### Performance and Capacity Specifications

Processing time for typical preclinical studies (20 animals, 10 timepoints) averages < 5 seconds on standard computers. Platform performance validation confirms handling of datasets up to the maximum specifications without degradation in user experience or computational accuracy.

#### Data Processing Workflow

The complete analytical workflow follows these automated steps:

1. <u>File Upload and Validation</u>: Structured CSV templates provided for data formatting, with automatic validation of required columns (Animal, Group, timepoint columns, optional Tumor_Weight)
2. <u>Automated Quality Control Assessment</u>: Automatic homogeneity evaluation and outlier detection using the Six-Criteria system
3. <u>Model Fitting and Quality Filtering</u>: Exponential parameter extraction with R^2^ quality filtering and validity assessment
4. <u>Analysis Selection</u>: User choice between complete analysis, animal-level filtering, or point-level filtering approaches
5. <u>Statistical Analysis</u>: Automated growth rate comparisons and TGR matrix generation with interactive exploration capabilities
6. <u>Results Export</u>: Enhanced data files and comprehensive HTML reports with publication-ready formatting

Data Export and Reproducibility Framework

#### Enhanced Data and Export Capabilities

ViVo generates augmented CSV files containing original measurements plus calculated parameters:

- Initial tumor burden (*N*_*0*_) with relative standard errors and confidence intervals
- Growth rates (*r*) and doubling times [ln(2)/r] for each valid animal
- R^2^ coefficients and exponential model quality metrics
- Outlier detection results with severity classifications and criteria explanations
- Predicted tumor weights for standardized timepoints when applicable

#### TGR Matrix Export Structure

Tumor Growth Rate matrices export as structured CSV format with the following column organization:

- Animal_ID: Unique animal identifier for traceability
- Experimental_Group: Treatment group designation for statistical comparisons
- *r*_*x*→*y*_ columns: All possible temporal interval growth rates as feature vectors

This standardized export format facilitates seamless integration with external statistical software and machine learning pipelines while maintaining complete analytical transparency.

#### Automated Report Generation

The platform generates comprehensive HTML reports that integrate:

- Raw data quality assessment with homogeneity scores and outlier detection summaries
- Exponential growth curve visualizations with model fit assessments
- Statistical results with publication-ready formatting including confidence intervals and effect sizes
- TGR matrices with statistical comparisons
- Complete methodology documentation for reproducibility

Reports include comprehensive analysis summaries suitable for results presentation or publication supplementary materials.

### Validation Design

#### Cross-Platform Validation and Performance Metrics

Primary validation used the publicly available Daskalakis dataset from subcutaneous xenograft studies. Additional validation encompassed four independent animal models across different cell line origins (human vs. murine), growth kinetics (rapid vs. indolent), implantation sites (mammary, pancreatic, subcutaneous), and immune status (athymic vs. immunocompetent). This systematic approach ensures methodological applicability beyond single experimental systems. Detailed model descriptions in Supplementary Note 3.

Performance assessment evaluated: **(1)** statistical power enhancement across sample sizes, **(2)** temporal pattern detection capability through TGR analysis, **(3)** tumor weight prediction accuracy for humane endpoint standardization, and **(4)** biological coherence validation through cross-platform kinetic consistency.

#### Animal ethics statement

All animal experiments were conducted in compliance with institutional guidelines and the European Directive 2010/63/EU for the protection of animals used for scientific purposes, and were approved by the Animal Welfare and Ethics Committee of University of Zaragoza Project Licence PI51/22 and PI68/23.

## Data and Code Availability

All source code for the ViVo platform is available at github.com/gcanudo-barreras/ViVo-Platform under the MIT License. The repository includes complete documentation, example datasets, and a live web application (gcanudo-barreras.github.io/ViVo-Platform/). Validation datasets from studies described in this work are included in the repository’s data folder. No restrictions apply to code access or usage.

## Acknowledgement

The authors thank projects PID2022-136861NB-I00, and PID2023-147471NB-I00 funded by MICIU/AEI/10.13039/501100011033, and project PDC2022-133376-I00, funded under the framework of the Recovery, Transformation and Resilience Plan, and financed by the European Union – NextGenerationEU, and Gobierno de Aragón (Research Group E07_23R) for financial support of our research. G. Canudo-Barreras also thanks Gobierno de Aragón for a predoctoral Grant. We thank Sandra Ardevines and Juan Carlos Morales-Solís for their critical review of the manuscript and honest insights, which greatly improved the quality of this work. We also thank Dra. Carmen Guerra for providing us with MLK 3287-Luc murine tumor cells.

## Author Contributions

G.C.B.: conceptualization, methodology, software, validation, formal analysis, investigation, data curation, writing, visualization. E.R.: resources, writing (review and editing), visualization. R.P.H.: conceptualization, writing (review and editing), visualization, project administration, funding acquisition. M.C.G.: conceptualization, supervision, writing (review and editing), visualization, project administration, funding acquisition.

