## Supplemental Data for "ViVo: A temporal modeling framework that boosts statistical power and minimizes animal usage"

### Supplementary Table S1: Exponential Fitting of the Daskalakis Dataset Data

| Animal ID | Group | $V_0$ (mm$^3$) | $r$ (day$^{-1}$) | $R^2$ * |
|---|---|---|---|---|
| 101 | Control | 55.2 | 0.208 | 0.9532 |
| 102 | Control | 111.8 | 0.137 | 0.8856 |
| 103 | Control | 29.1 | 0.270 | 0.9525 |
| 104 | Control | 46.0 | 0.211 | 0.9663 |
| 105 | Control | 27.8 | 0.309 | 0.9516 |
| 106 | Control | 42.4 | 0.200 | 0.9625 |
| 107 | Control | 54.3 | 0.358 | 0.9524 |
| 108 | Control | 60.5 | 0.365 | 0.9276 |
| 201 | Drug | 49.7 | 0.183 | 0.9735 |
| 202 | Drug | 63.9 | 0.183 | 0.9075 |
| 203 | Drug | 23.8 | 0.161 | 0.8981 |
| 204 | Drug | 28.6 | 0.242 | 0.9376 |
| 205 | Drug | 36.6 | 0.186 | 0.9775 |
| 206 | Drug | 35.8 | 0.114 | 0.8088 |
| 207 | Drug | 43.9 | 0.039 | **0.4700** |
| 208 | Drug | 47.5 | 0.203 | 0.9407 |

| | | | | |
|---|---|---|---|---|
| 209 | Drug | 38.9 | 0.160 | 0.9368 |
| 210 | Drug | 108.2 | 0.144 | 0.8688 |

* Animals that do not meet the $R^2$ quality criteria are excluded. $R^2 < 0.8$ (red), $R^2 < 0.9$ (blue).

**Supplementary Table S2: Daily Caliper Comparisons of the Daskalakis Dataset Data (Raw Statistical Analysis)**

| Day | Control mean | Drug mean | p-value | Cohen’s d | Significance*[1] |
|---|---|---|---|---|---|
| 0 | 55.6 | 51.8 | 0.5338 | 0.323 | ns |
| 3 | 97.0 | 63.9 | 0.0205 | 1.089 | * |
| 4 | 146.8 | 75.1 | 0.0266 | 1.081 | * |
| 5 | 195.2 | 104.1 | 0.0343 | 0.907 | * |
| 6 | 251.2 | 141.4 | 0.1728 | 0.818 | ns |
| 7 | 550.6 | 217.3 | 0.0831 | 0.859 | ns |
| 10 | 832.9 | 269.9 | 0.0266 | 1.273 | * |
| 11 | 1041.5 | 323.8 | 0.0021 | 1.300 | ** |
| 12 | 754.8 | 413.0 | 0.0934 | 1.177 | ns |
| 13 | 911.5 | 473.0 | 0.0727 | 1.244 | ns |
| 14 | 1330.7 | 696.8 | 0.0420 | 1.283 | * |
| 17 | 1528.3 | 788.9 | 0.0400 | 1.506 | * |
| 18 | 1692.1 | 884.6 | 0.0240 | 1.678 | * |

*[1] Statistical significance assessed using Mann-Whitney U tests with Cohen's d effect size calculations. Significance levels: *p<0.05, **p<0.01, ***p<0.001.

**Supplementary Table S3: Daily BLI Comparisons of the Orthotopic 4T-1-Luc2 Assay (Raw Statistical Analysis)**

| Day | Control mean | Drug mean | p-value | Cohen’s d | Significance*[1] |
|---|---|---|---|---|---|
| 0 | 5.74E+08 | 3.75E+08 | 0.2786 | 0.703 | ns |
| 2 | 5.70E+08 | 6.45E+08 | 1 | -0.204 | ns |

| | | | | | |
|---|---|---|---|---|---|
| 4 | 4.42E+09 | 2.54E+09 | 0.0148 | 1.145 | * |
| 7 | 4.03E+10 | 1.15E+10 | 0.0003 | 2.681 | *** |

*[1] Statistical significance assessed using Mann-Whitney U tests with Cohen's d effect size calculations. Significance levels: *p<0.05, **p<0.01, ***p<0.001.

### Supplementary Table S4: Daily Caliper Comparisons of the Xenograft 4T-1-Luc2 Assay (Raw Statistical Analysis)

| Day | Control mean | Drug mean | p-value | Cohen's d | Significance*[1] |
|---|---|---|---|---|---|
| 11 | 199.1 | 227.8 | 0.3416 | -0.266 | ns |
| 14 | 348.8 | 307.7 | 0.9159 | 0.232 | ns |
| 18 | 533.4 | 346.6 | 0.4595 | 0.665 | ns |
| 21 | 832.1 | 403.9 | 0.0447 | 1.194 | * |
| 25 | 1089.0 | 524.2 | 0.0527 | 1.194 | ns |
| 32 | 1742.8 | 826.1 | 0.0049 | 1.381 | ** |
| 39*[2] | - | - | - | - | - |

*[1] Statistical significance assessed using Mann-Whitney U tests with Cohen's d effect size calculations. Significance levels: *p<0.05, **p<0.01, ***p<0.001. *[2] No comparison was possible because the control group consisted of only one animal.

### Supplementary Table S5: Daily Caliper Comparisons of the MDA-MB-231-luc2-GFP Assay (Raw Statistical Analysis)

| Day | Control mean | Drug mean | p-value | Cohen's d | Significance*[1] |
|---|---|---|---|---|---|
| 22 | 46.7 | 80.8 | 0.4225 | -0.822 | ns |
| 25 | 102.3 | 147.0 | 0.4225 | -0.580 | ns |
| 29 | 149.3 | 227.1 | 0.3939 | -0.644 | ns |
| 32 | 194.2 | 356.9 | 0.2403 | -0.870 | ns |

*[1] Statistical significance assessed using Mann-Whitney U tests with Cohen's d effect size calculations. Significance levels: *p<0.05, **p<0.01, ***p<0.001.

### Supplementary Table S6: Daily BLI Comparisons of the MLK 3287-Luc Assay (Raw Statistical Analysis)

| Day | Control mean | Drug mean | p-value | Cohen's d | Significance*[1] |
|---|---|---|---|---|---|
| 3 | 1.32E+08 | 1.32E+08 | 0.6232 | 0.000 | ns |
| 7 | 6.91E+08 | 4.50E+08 | 0.5708 | 0.410 | ns |
| 10 | 1.03E+09 | 6.93E+08 | 0.2123 | 0.572 | ns |
| 14 | 1.31E+09 | 8.01E+08 | 0.1620 | 0.748 | ns |

*[1] Statistical significance assessed using Mann-Whitney U tests with Cohen's d effect size calculations. Significance levels: *p<0.05, **p<0.01, ***p<0.001.

### Supplementary Table S7: TGR Intervals Comparisons of the Xenograft 4T-1-Luc2

| $Day_x \rightarrow day_y$ | Control mean (n) | Drug mean (n) | p-value | Cohen's d | Significance*[1] |
|---|---|---|---|---|---|
| 11→14 | 0.168 (10) | 0.102 (8) | 0.004 | 1.498 | ** |
| 11→18 | 0.136 (10) | 0.056 (8) | 0.003 | 2.359 | ** |
| 11→21 | 0.142 (10) | 0.060 (8) | < 0.001 | 3.546 | *** |
| 11→25 | 0.124 (10) | 0.060 (8) | 0.001 | 2.903 | ** |
| 11→32 | 0.111 (9) | 0.052 (8) | 0.001 | 2.334 | ** |
| 11→39 | 0.062 (1)*[2] | 0.0523 (4) | - | - | - |
| 14→18 | 0.084 (10) | 0.033 (8) | 0.006 | 1.510 | ** |
| 14→21 | 0.122 (10) | 0.039 (8) | < 0.001 | 3.269 | *** |
| 14→25 | 0.102 (10) | 0.055 (8) | < 0.001 | 2.791 | *** |
| 14→32 | 0.102 (9) | 0.048 (8) | 0.004 | 1.858 | ** |
| 14→39 | 0.064 (1)*[2] | 0.047 (4) | - | - | - |
| 18→21 | 0.158 (10) | 0.053 (8) | < 0.001 | 2.432 | *** |
| 18→25 | 0.118 (10) | 0.056 (8) | 0.004 | 1.972 | ** |
| 18→32 | 0.099 (9) | 0.046 (8) | 0.009 | 1.534 | ** |
| 18→39 | 0.066 (1)*[2] | 0.0512 (4) | - | - | - |

| | | | | | |
|---|---|---|---|---|---|
| 21→25 | 0.071 (10) | 0.057 (8) | 0.722 | 0.168 | ns |
| 21→32 | 0.077 (9) | 0.047 (8) | 0.149 | 0.662 | ns |
| 21→39 | 0.054 (1)*[2] | 0.045 (4) | - | - | - |
| 25→32 | 0.0796 (9) | 0.0496 (8) | 0.102 | 0.785 | ns |
| 25→39 | 0.064 (1)*[2] | 0.044 (4) | - | - | - |
| 32→39 | 0.048 (1)*[2] | 0.026 (4) | - | - | - |

*[1] Statistical significance assessed using Mann-Whitney U tests with Cohen's d effect size calculations. Significance levels: *p<0.05, **p<0.01, ***p<0.001. *[2] No possible statistical comparison due to insufficient sample size.

### Supplementary Table S8: Parameters used to compute the homogeneity score

| Parameter | Symbol | Default value | Interpretation |
|---|---|---|---|
| Excellent threshold | ε | 15 | CV at or below this value receives maximal base score (100) |
| Good threshold | *Good* | 25 | Upper bound of the “good” interval (used for categorical labels) |
| Poor threshold | π | 30 | CV above this value receives an accelerated penalty (slope = −2) |
| Small-sample factor (applies if n<5) | $S_{small}$ | 0.8 | Multiplies the base score for small groups (20% reduction) |
| Very-small-sample factor (applies if n<3) | $S_{verySmall}$ | 0.6 | Additional multiplier for very small groups (40% reduction) |

### Supplementary Table S9: Adaptative configurations for the Six-Criteria Outlier Detector based on the sensitivity levels.

| Sensitivity Level | Sample size | Max Growth (/day) | Max Decline (/day) | IQR Sensitivity | Notes |
|---|---|---|---|---|---|
| **Ultra-Conservative** | n < 5 | 5000% | 90% | 4.0 | Requires multiple indicators for outlier detection |
| **Conservative** | n = 5-10 | 2000% | 80% | 3.0 | Requires multiple indicators for outlier detection |

| | | | | | |
|---|---|---|---|---|---|
| **Moderate** | n > 10 | 1000% | 67% | 2.0 | One indicator is sufficient for outlier detection |

### Supplementary Note 1: Error Propagation Analysis for Exponential Growth Models

#### Mathematical Framework

For an exponential growth function of the form: $N(t) = N_0 e^{rt}$

where $N_0$ is the initial tumor burden, $r$ is the growth rate, and $t$ is time, we derive the propagation of relative standard errors (RSE) from the model parameters to the predicted values.

#### Mathematical Derivation

Taking the natural logarithm of both sides:

$$\ln N(t) = \ln N_0 + rt$$

Differentiating with respect to the parameters:

$$dN / N = dN_0 / N_0 + t dr$$

This yields the relative error as:

$$\delta N / N = \delta N_0 / N_0 + t \delta r$$

where δ$N$, δ$N_0$, and δ$r$ represent the absolute errors in $N(t)$, $N_0$, and $r$, respectively.

Squaring both sides:

$$\left(\delta N / N\right)^2 = \left(\delta N_0 / N_0\right)^2 + 2t(\delta N_0 / N_0)\left(\delta r\right) + t^2\left(\delta r\right)^2$$

Assuming independence between parameter estimation errors (i.e., $E[(\delta N_0/N_0)(\delta r)] = 0$), the expected value becomes:

$$E\left[\left(\delta N / N\right)^2\right] = E\left[\left(\delta N_0 / N_0\right)^2\right] + t^2 E\left[\left(\delta r\right)^2\right]$$

#### Result

The relative standard error of the predicted tumor size is given by:

$$RSE\left(N\right) = \sqrt{\left[RSE^2\left(N_0\right) + t^2 \cdot RSE^2\left(r\right)\right]}$$

#### Theoretical Implications

This decomposition reveals that:

1. **Direct contribution**: Uncertainty in the initial size $N_0$ contributes directly to prediction uncertainty.
2. **Quadratic amplification**: Uncertainty in the growth rate ($r$) contributes quadratically with time, becoming the dominant source of error for extended studies.

3. **Variance reduction mechanism**: When comparing growth rates between treatment groups, the $N_0$ terms largely cancel out, leaving primarily the more precisely estimated $r$ parameters.

**Validation Against Empirical Data**

Using the Daskalakis dataset, we validated this theoretical framework:

- **Predicted RSE reduction**: Formula suggests RSE($r$) < RSE($V_0$) for multi-timepoint data.
- **Empirical confirmation**: Observed RSE($r$)/RSE($V_0$) ratios of 0.46-0.56 consistent with theoretical predictions.
- **Time dependency**: Longer observation periods show greater variance reduction, as predicted by $t^2$ coefficient.

**Practical Applications**

This error propagation formula enables:

1. **Quantitative assessment** of prediction confidence intervals.
2. **Parameter importance ranking** identifies which parameter contributes most to model uncertainty.
3. **Experimental design optimization** suggests that precise estimation of tumor growth rate ($r$) is particularly critical for extended studies.
4. **Sample size planning** enables a priori estimation of expected RSE ratios for power calculations.

**Connection to Statistical Power Enhancement**

The mathematical framework explains why exponential modeling consistently outperforms size-based comparisons:

- **Volume/BLI comparisons**: Subject to full measurement error at each timepoint.
- **Growth rate comparisons**: Benefit from error averaging across multiple timepoints.
- **Temporal integration**: Mathematical constraints of exponential form reduce effective noise.
- **Between-group comparisons**: $N_0$ uncertainties largely cancel, emphasizing biologically relevant $r$ differences.

This theoretical foundation validates the empirical power enhancement observed across all tested sample sizes and provides a quantitative framework for predicting method performance in future studies.

### Supplementary Note 2: Cohen's d Effect Size Interpretation

Cohen's d quantifies the standardized difference between two group means, providing a scale-independent measure of effect magnitude that complements p-value significance testing. In our tumor growth rate analysis, Cohen's d was calculated as:

$$d = \frac{(mean_1 - mean_2)}{Sp}$$

Where,

$$\text{Sp} = \sqrt{\left[\frac{\left((n_1-1)\times SD_1^2 + (n_2-1)\times SD_2^2\right)}{(n_1+n_2-2)}\right]}$$

Interpretation guidelines following Cohen's conventions:

- Small effect: d = 0.2-0.5 (subtle but meaningful difference)
- Medium effect: d = 0.5-0.8 (moderate practical significance)
- Large effect: d ≥ 0.8 (substantial practical significance)

In clinical contexts, effect sizes ≥0.8 indicate therapeutically relevant differences that are likely to translate into meaningful clinical outcomes. Our TGR analysis revealed predominantly large effect sizes (d > 1.0), indicating robust drug efficacy that extends beyond statistical significance to practical therapeutic relevance.

Cohen's d is particularly valuable for tumor growth studies because:

1. **Scale independence**: Allows comparison across different measurement units and timepoints.
2. **Clinical relevance**: Large effect sizes (d ≥ 0.8) suggest clinically meaningful tumor growth inhibition.
3. **Power analysis**: Facilitates sample size calculations for future studies.
4. **Meta-analysis compatibility**: Enables integration with broader literature syntheses.

The consistently large effect sizes observed across our therapeutic windows (d = 1.105-1.583) demonstrate not only statistical significance but substantial biological and potential clinical impact.

### Supplementary Note 3: Mouse Models

#### Syngeneic Orthotopic Model with 4T-1-Luc2 Cells

A syngeneic orthotopic murine breast cancer model was established by implanting 4T-1-Luc2 cells (CRL-2539-LUC2™ / ATCC) into the mammary fat pad of athymic nude mice (Rj:ATHYM-*Foxn1*$^{nu}$-Janvier), which are maintained on a BALB/cAnNRj genetic background. Despite their immunodeficiency, these mice share sufficient genetic compatibility with the 4T-1 cell line to support a syngeneic tumor environment. The model was used to evaluate the efficacy of a compound during the initial 10 days of treatment (days 0, 2, 4, 7 and 10). Tumor progression was monitored by bioluminescence imaging (BLI). Mice were randomly assigned to two experimental groups: control and treatment (n=8,8).

#### Xenogeneic Orthotopic Model with MDA-MB-231-luc2-GFP Cells

A xenogeneic orthotopic model of human triple-negative breast cancer was established by implanting MDA-MB-231-luc2-GFP cells (Caliper Life Sciences) into the mammary fat pad of athymic nude mice. The model was used to evaluate the efficacy of a compound over a 32-day period. Tumor resection was performed on day 22 to assess metastatic progression. Tumor volume was estimated using caliper (Equation S1). The study included two experimental groups: control and treatment (n=6,6).

**Orthotopic Model with Syngeneic-like MLK 3287-Luc Cells**

An orthotopic murine pancreatic ductal adenocarcinoma (PDAC) model was established by implanting syngeneic-like MLK 3287-Luc cells into athymic nude mice to evaluate the efficacy of a compound over a 14-day treatment period. Tumor growth was measured by bioluminescence imaging (BLI). Animals were divided into two experimental groups: control and treatment (n=8,8).

The MLK 3287-Luc murine tumor-derived cell line was established in the laboratory of Dr. Carmen Guerra (https://orcid.org/0000-0002-3891-046X) from a genetically engineered mouse model generated by crossing multiple transgenic mouse lines carrying the following alleles: *ELA* (+/E), *TetO-Cre* (+/T), *Kras* (+/LSL-G12Vgeo), *p53* (lox/lox), *EGFR* (+/+), *ROSA26-LSL-YFP* (KI/KI), and *TgpCAG-LSL-KFP* (+/+).

Although athymic nude mice are immunodeficient, the murine origin and background of the implanted cells ensure a genetically compatible tumor microenvironment, making this model functionally syngeneic.

**Subcutaneous Syngeneic Model with 4T-1-Luc2 Cells**

A subcutaneous syngeneic murine breast cancer model was established by injecting 4T-1-Luc2 cells (CRL-2539-LUC2™ / ATCC) into immunocompetent mice (BALB/cByJRj-Janvier). The model was used to evaluate the efficacy of a compound from day 11 to day 39 of treatment (days 11, 14, 18, 21, 25, 32 and 39). Tumor volume was estimated using caliper (Equation S1). Mice were randomized into two groups: control and treatment (n=10,11).
